# A measure of the somatosensory blink reflex to parametric variation of mechanical stimulation

**DOI:** 10.1101/2022.07.28.501855

**Authors:** Eric A Kaiser, Edda B Haggery, Dena P Garner, Vatinee Y Bunya, Geoffrey K Aguirre

## Abstract

The primary goal of this study was to develop a parametric model that relates variation in stimulation of the trigeminal nerve to properties of the blink response. We measured blink responses in 17 healthy, adult participants to air puffs directed at the lateral canthus of the eye at five different, log-spaced intensities (3.5 - 60 PSI). Lid position over time was decomposed into amplitude and velocity components. We found that blink amplitude was systematically related to log stimulus intensity, with the relationship well described by a sigmoidal function. The parameters of the model fit correspond to the slope of the function and the stimulus intensity required to produce half of a maximal blink response (the half-response threshold). There was a reliable decrease in the half-response threshold for the contralateral as compared to the ipsilateral blink response. This decrease was consistent across participants despite substantial individual differences in the half-response threshold and slope parameters of the overall sensitivity function, suggesting that the laterality effect arises in the neural circuit subsequent to individual differences in sensitivity. Overall, we find that graded mechanical stimulation of the somatosensory trigeminal afferents elicits a graded response that is well described by a simple parametric model. We discuss the application of parametric measurements of the blink response to the detection of group differences in trigeminal sensitivity.

**Impact statement:**

- There has been limited study of how variation in stimulus intensity influences the cutaneous blink response. We describe how variation in the intensity of an air-puff stimulus relates to the amplitude of the blink response using a two-parameter model. The parameters of the model capture individual differences in response, which may be used in future studies to identify population differences in trigeminal sensitivity.

## 1. Introduction

The somatosensory blink reflex is an easily evoked, clinically relevant measure of brainstem function. The cornea of the eye and periorbital skin are innervated by the ophthalmic division (V1) of the fifth cranial nerve (CN V), also known as the trigeminal nerve. In the brainstem, the principal sensory nucleus of V receives light-touch sensation from V1, while the nucleus caudalis receives both light touch and nociceptive afferents (May & Porter, 1998). Both of these nuclei project to local brainstem circuits that serve reflexive functions, including eye blink (May & Warren, 2021).

There is a long history of quantitative measurement of the human blink reflex elicited by a somatic stimulus, with a frequent approach being unilateral electrical stimulation of the supraorbital nerve or cornea, and electromyographic (EMG) measurement of the response. This traditional approach assesses isolated components at the expense of some ecological validity. Specifically, the somatosensory receptive surface is bypassed in favor of direct electrical stimulation of the nerve, and the isolated electrical response of the orbicularis oculi muscle is measured as opposed to the movement of the lid (which is influenced by three muscles and the elastic properties of the tissue). One goal of the current study was to obtain a measure of the integrated blink reflex to somatosensory stimulation.

In usual practice, the evoked response is measured for a fixed stimulus intensity, or the threshold stimulus is found that evokes a criterion EMG response. There has been relatively little study of the effect of variation in somatosensory stimulus intensity. A focus upon measurement at response threshold can obscure individual and population differences. For example, two groups may initiate a physiologic response at the same threshold but differ in the amplification of that response with further increases in stimulus intensity. Examination of the sensitivity function that relates stimulus to response, and how this function interacts with other experimental variables, may be revealing as to the organization of the underlying neural system.

In the rabbit, a graded change in orbicularis EMG response is found in response to variation in the intensity of an air puff stimulus (Gruart, Schreurs, del Toro, & Delgado-Garcia, 2000). There are suggestions of a similar effect in human data. In a review of the clinical assessment of the blink reflex, Esteban informally observed that increases in the electrical current used to stimulate the supraorbital nerve evoked larger and earlier EMG components from the orbicularis oculi (Esteban, 1999). Flaten and Blumenthal found that varying the intensity of an air puff (directed at the temple) between 1.5 and 4.5 PSI resulted in a monotonic increase in the amplitude of the EMG voltage response from the orbicularis (Flaten & Blumenthal, 1998). A second goal of the current study was to more fully characterize how the blink reflex varies with graded changes in somatic stimulation.

We report here measurements of the somatosensory blink reflex evoked by mechanical stimulation of varying intensity. Specifically, we directed an air puff stimulus to the lateral canthus of the eye at logarithmically spaced pressure levels and measured the change in lid position with high-speed infrared (IR) cameras. We approached the data within a modeling framework that extracts the amplitude and latency of the blink and characterizes the sensitivity function that relates blink response to variation in stimulus intensity. To anticipate our results, we find reliable differences in the sensitivity functions of the ipsilateral and contralateral response, individual differences in the sensitivity of the somatosensory blink reflex, and changes in response across trials as a measure of habituation.

## 2. Methods

### 2.1 Participants

We studied 17 participants recruited from The Citadel in Charleston, SC (a public military college) and from non-clinical settings in the surrounding Charleston community. Participants were at least 18 years of age, had no history of concussion within the past 5 years, and denied a history of recurrent headache. Nine of the subjects identified as women, and the mean age of all participants was 25 years (SD ± 4.0; range 20 to 32). This study was approved by The Citadel Institutional Review Board, and all participants provided written informed consent. All experiments were performed in accordance with relevant guidelines and regulations.

Our pre-registered enrollment goal was 15 subjects, and 17 were studied as the data from 2 subjects did not meet our original inclusion criteria. The analysis approach presented here departs substantially from our pre-registered plan and allowed us to retain the measurements from all subjects. The results of the original planned analyses can be found with the pre-registration document: https://github.com/gkaguirrelab/preregistrations/tree/master/blink_2021.

### 2.2 Operation of the device

Blink responses were evoked and measured using a custom-modified, FDA cleared device (The EyeStat device, Gen3, Blink TBI Inc). The device was positioned over the eyes of the seated subject. Air puffs were produced by high-purity (“food grade”) CO_2_ cannisters, minimizing the potential for introducing noxious solid particles to the cornea, and administered at pressures below those used in ocular tonometry. During each of many ∼30 second acquisitions, the device directed 8, temporally spaced, 100 ms air puffs (i.e., 8 “trials”) of a given pressure level. The air puff was centered on the lateral canthus of the eye, with a radius of ∼5.5 mm (Figure S1a). The laterality (left or right eye) of each trial was selected at random. The inter-stimulus-interval was jittered to reduce the ability of the participant to anticipate the timing of the stimulus, with at least 3 seconds between trials. The eyes were illuminated with a ring of IR light emitting diodes that produced a corresponding ring of glint reflections on the tear film. A pair of high-speed IR cameras (300 Hz, 800 x 600 pixels) were used to measure the timing of obscuration of the glints by the movement of the eyelids. Recordings were made of the movement of the eyelid ipsilateral and contralateral to the side on which the stimulus was delivered. Microphones adjacent to the lateral canthus detected the arrival time of the air puff, and the timing of the pressure wave was used to synchronize recorded responses across trials.

During testing, subjects steadied themselves to receive the stimulus by holding the legs of a tripod that supported the device. Room lights were left on during the session, although the field of view of the subject was dark during data acquisition while their face was placed within the device. The stimulus was accompanied by an audible hiss; we measured the sound pressure amplitude and temporal profile of the auditory component of the stimulus (Figures S1b, c).

The device used in this study was modified by the manufacturer to externalize pressure regulation. The CO_2_ cartridge was connected by rigid tubing to an external pressure regulator (Reg3000x, Leland Gas Technologies, South Plainfield, NJ), and then returned by tubing to the device. Accuracy of specified air pressure levels was monitored with a National Institute of Standards and Technology calibrated device (PressureMon2000, RegCo, Miami Florida), which is certified to be accurate within ±0.2 pounds per square inch (PSI; 1 PSI = ∼6.9 kilopascals).

### 2.3 Data collection

The stimulus intensity was varied in separate acquisitions between five different, log-spaced pressure values. A set of 26 acquisitions (each composed of 8 air puff trials of the same stimulus pressure) were obtained in each of two sessions, with the order of pressure values following a pseudo-random, counterbalanced, de Bruijn sequence (Aguirre, Mattar, & Magis-Weinberg, 2011): [3, 3, 0, 2, 4, 4, 2, 0, 1, 3, 1, 1, 0, 4, 3, 2, 2, 3, 4, 1, 2, 1, 4, 0, 0, 3]; where 0 = 3.5 PSI, 1 = 7.5 PSI, 2 = 15 PSI, 3 = 30 PSI, and 4 = 60 PSI. We discarded the data from all trials from the first acquisition in each session, thus producing a set 25 acquisitions in each session that provided full counterbalance of stimulus levels. Further, in those analyses that were not designed to examine the effect of habituation across trials, we did not include the response to the first trial in each 8-trial acquisition. Thus, a total of 10 acquisitions were obtained at each of the five pressure levels, with each acquisition containing seven retained trials. The operator of the device attempted to achieve the desired PSI value within ±1 PSI, and the actual, achieved stimulus intensity was recorded for each acquisition. The unsigned mean deviation from nominal values was 0.31 PSI, and in all cases did not exceed 1 PSI. For one participant, 3 of the 50 acquisitions included 4 trials instead of 8 due to operator error; we elected to retain the data from this subject as this change in trial number was expected to have a negligible effect upon the data.

There was a 30 – 60 second break following each acquisition. Each session was approximately 50 minutes in duration, and the two sessions were separated by 1 to 5 days.

### 2.4 Data pre-processing

The raw data consisted of IR videos of the eyelids during each trial. Initial processing of these videos was conducted by Blink TBI Inc. using proprietary software. This first stage of analysis provided a measure of lid position during each trial with 3.33 ms temporal resolution, with all trials aligned to the time of arrival of the stimulus at the eye. All subsequent analyses were performed using custom MATLAB software (Mathworks, Natick, Massachusetts).

The lid position over time was obtained for each subject for each stimulus intensity level, averaging over all acquisitions and trials. Because spatial calibration of the camera was not performed for each participant, we did not have a measurement of the width of the palpebral fissure in absolute units. Instead, the raw measurement of lid position in units of pixels was converted to a proportion of full eye closure by scaling all responses by the maximum excursion observed for that subject and session. Full lid closure was observed for all subjects at either the 30 or 60 PSI stimulus. This normalization accounted for variation in the absolute size of the palpebral fissure across subjects and in the precise distance of the eye from the IR cameras.

The primary analyses of the study examined the set of average lid position vectors obtained from the 17 subjects at each of the 5 pressure levels, measured from the eye ipsilateral and contralateral to the stimulus, averaged across trials and acquisitions.

### 2.5 Modeling and summary measures of the blink response

An initial summary of the data was provided by measuring the amplitude, latency, and duration of the across-subject average blink response as a function of stimulus intensity. The amplitude was expressed as the proportion of lid closure. Latency was expressed as the time after stimulus arrival by which the lid had closed by 10%. Duration was expressed as the time period during which the lid was positioned at more than half of the maximum closure. The standard error of these measures was obtained by bootstrap resampling across subjects.

A model of the blink response over time was created to allow the temporal vector of lid position to be summarized with a small set of coefficients. The first component of the model was the average profile of lid position over time, combined across subjects, pressure levels, and the ipsilateral and contralateral eye. The second component was the first derivative of average lid position. This component reflects the velocity of the blink and could be used to calculate the latency of the blink response. Next, a set of residual vectors was obtained by regressing each of the 170 measures (17 subjects x 2 eyes x 5 stimulus levels) of lid position upon these first two components. The set of residuals was then subject to a principal components analysis, and the first eigenvector was retained as the third component of the model of temporal response.

A simple linear regression was then used to fit this three-component model to each lid-position vector, yielding three coefficients. The first coefficient was expressed in units of proportion of maximal lid closure. We converted the values for the 2^nd^ coefficient from amplitude to latency (ms) by calculating the effect of this coefficient upon the timing of the modeled response. The values for the 3^rd^ coefficient were left as arbitrary amplitude units. The effect of stimulus intensity upon the coefficients was then examined, as well as the effect of other variables. This fitting approach leverages the entire profile of the blink response for estimation of amplitude and latency. Because the lid position vector is smooth in time, our analysis is able to estimate latency at a temporal resolution higher than the sampling rate of the measurement (Henson, Price, Rugg, Turner, & Friston, 2002).

### 2.6 Psychometric function relating blink amplitude and stimulus pressure

A Weibull cumulative distribution function (Wichmann & Hill, 2001) was used to model the effect of log-transformed stimulus pressure upon the amplitude of blink response. Two parameters controlled the shape of the fit, corresponding to the maximum slope of the function and the pressure at which 50% eye closure was predicted. This function was fit to the data from each subject using the MATLAB *fmincon* function. A comparison of the fit parameters between the ipsilateral and contralateral eye was performed with a paired t-test across subjects. Bootstrap resampling across acquisitions was used to estimate the standard error of the mean of the parameters within subjects.

### 2.7 Examination of habituation effects

For each subject, the blink response vectors for sequential trials within each acquisition were obtained and averaged over stimulus levels. After fitting with the three-component temporal model, the coefficients were examined for an effect of trial number. For this analysis, all 8 trials in each acquisition were retained (i.e., the first trial in each acquisition was included).

### 2.8 Examination of auditory component of the air puff stimulus

See supplementary materials.

## 3. Results

### 3.1 Blink amplitude and velocity increase with stimulus intensity

We measured the somatosensory blink reflex in 17 participants in response to an air puff stimulus (Figure 1a). A total of 10 acquisitions of 8 air puff trials were obtained at each of five, log-spaced pressure intensities, with the order of acquisitions following a counter-balanced sequence (Figure 1b). IR video recordings of the eye ipsilateral and contralateral to the stimulus measured lid position over time (Figure 1c). From these recordings, the average blink response was obtained for each participant for each of the five stimulus intensities.

**Figure 1.**
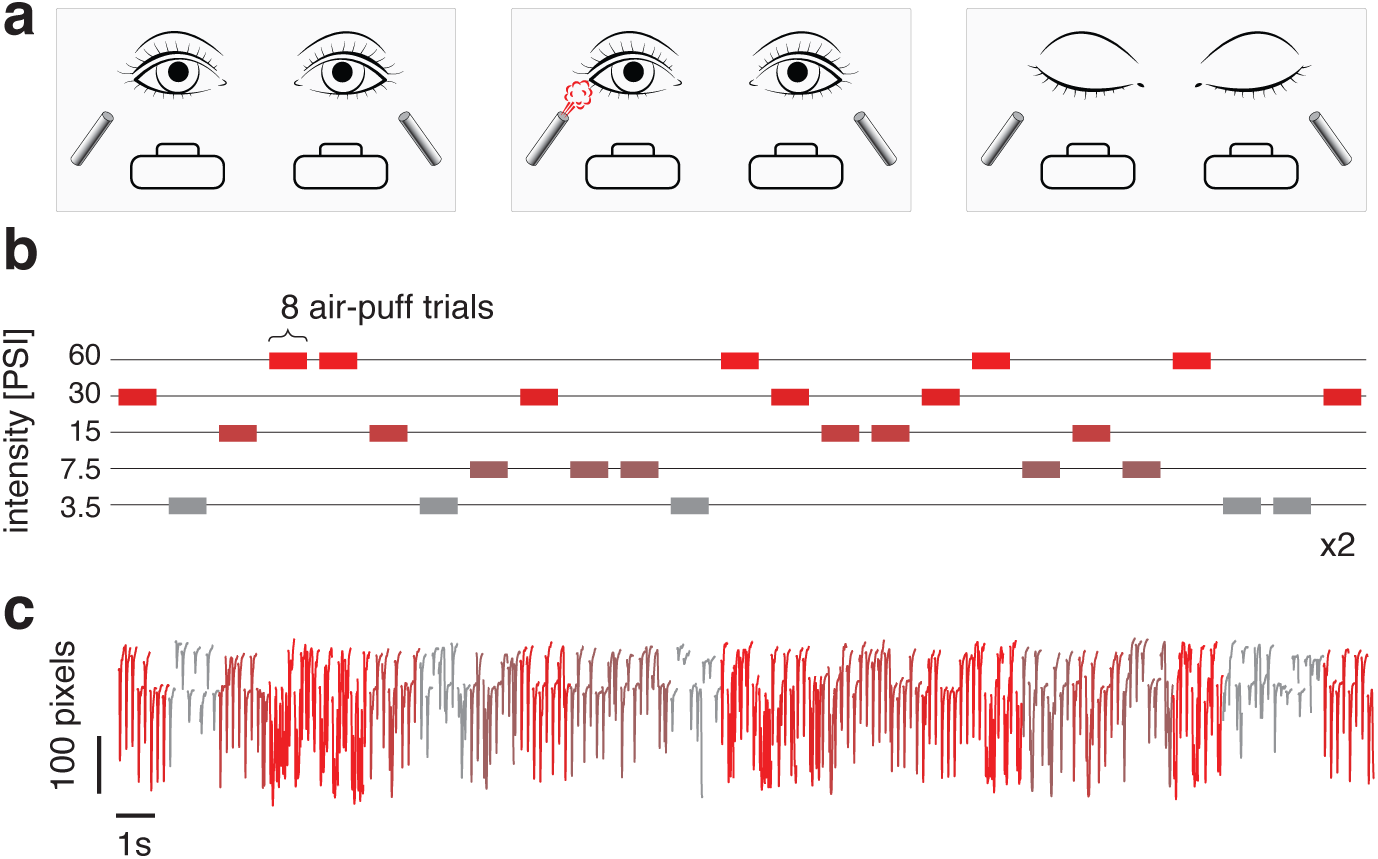
Experiment overview. **(a)** Blink responses were measured in response to cutaneous stimulation. The participant placed their face within a device that positioned air delivery tubes adjacent to the lateral canthi. At unpredictable intervals an air puff was delivered to the left or right eye. A pair of high-speed IR cameras recorded the resulting movement of the eyelids. **(b)** Counterbalanced pressure sequence. Each dash represents a single acquisition that presented a sequence of 8 air puff trials. The air pressure used for the 8 trials was varied across acquisitions as shown. The entire sequence was administered twice. **(c)** Lid position over time obtained for an example subject. The data from each acquisition is color coded to indicate stimulus intensity, following the convention shown in panel b.

We first combined the responses from the eyelids ipsilateral and contralateral to the stimulus and examined the evoked blink response across participants and stimulus intensity levels. Figure 2a presents these data for each subject as a temporal raster plot. Each evoked response is in a row with the grayscale level indicating the amplitude of lid movement. A similar general pattern was seen across subjects (grouped blocks) and stimulus intensity levels (increasing stimulus intensity across rows within a block): Following stimulus arrival (vertical blue line), a blink of 100-200 ms duration was initiated after a latency of 40-60 ms. Examination of the average responses across subjects (Figure 2b and Table 1) demonstrates that increasing stimulus intensity evokes a blink response of larger amplitude and a shorter latency.

**Figure 2.**
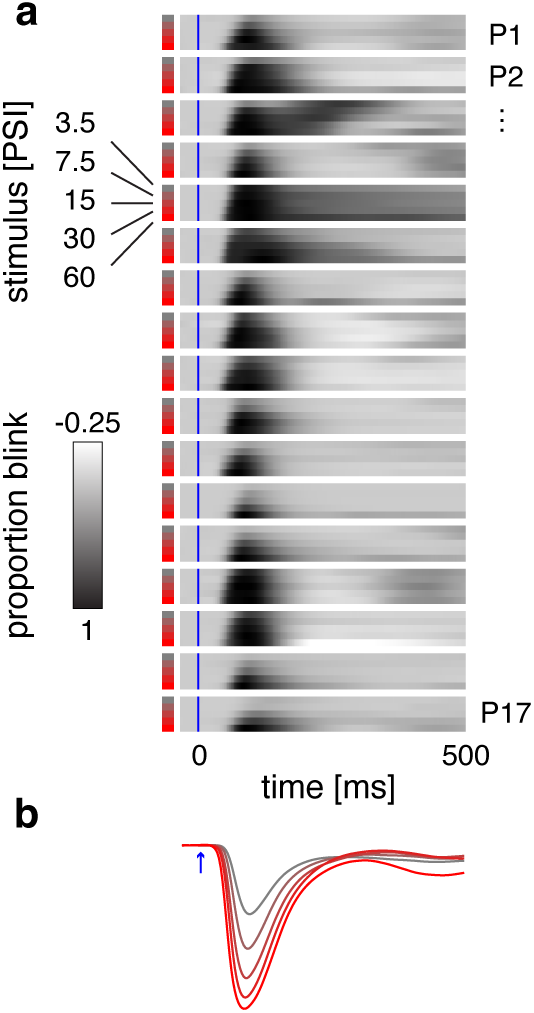
Observed blink responses. **(a)** The raster plot presents the temporal profile of lid position (averaged across acquisitions and ipsilateral and contralateral eyes) on each row, with blink amplitude indicated by grayscale shading. The responses to each of the five stimulus intensity levels are grouped together with each block running vertically. Each block corresponds to a different participant (P1, P2, etc). The time of stimulus arrival at the cornea is indicated by a blue line. **(b)** The average (across subject) blink response evoked by the five stimulus intensity levels (color coding following the convention in Figure 1b).

**Table 1.** Summary measures of average blink response. The properties of the mean (±SEM across subject) blink response evoked by varying levels of mechanical stimulation. Proportion closure is the degree to which the lid achieved the maximum possible closed position. The latency is the time to reach 10% eye closure following the arrival of the stimulus. The duration is the time during which the lid was at greater than 50% of maximum closure.

| Stimulus intensity (PSI) | Proportion closure | Latency (ms) | Duration (ms) |
| --- | --- | --- | --- |
| 3.5 | 0.40 ± 0.1 | 61.1 ± 2.8 | 86.8 ± 23.5 |
| 7.5 | 0.61 ± 0.1 | 54.5 ± 1.8 | 83.8 ± 6.1 |
| 15.0 | 0.78 ± 0.1 | 49.2 ± 1.9 | 90.4 ± 6.2 |
| 30.0 | 0.90 ± 0.0 | 45.2 ± 1.7 | 96.1 ± 7.2 |
| 60.0 | 0.97 ± 0.0 | 40.2 ± 1.1 | 106.4 ± 7.5 |

Our initial summary measures (Table 1) were derived from single time points in the blink response. We developed a temporal model that leverages the entire temporal profile of lid position to improve the precision and reliability of our derived measures. The model was composed of three orthogonal, smooth temporal components (Figure 3a), the first two of which describe variation in the amplitude and velocity of the evoked response. For a given blink response, we derived the coefficients that best scale the components of the model to fit the data (Figure 3b). We fit this model to the blink response from each participant at each stimulus level and found that it provided a good description of the data, explaining over 95% of the variance.

**Figure 3.**
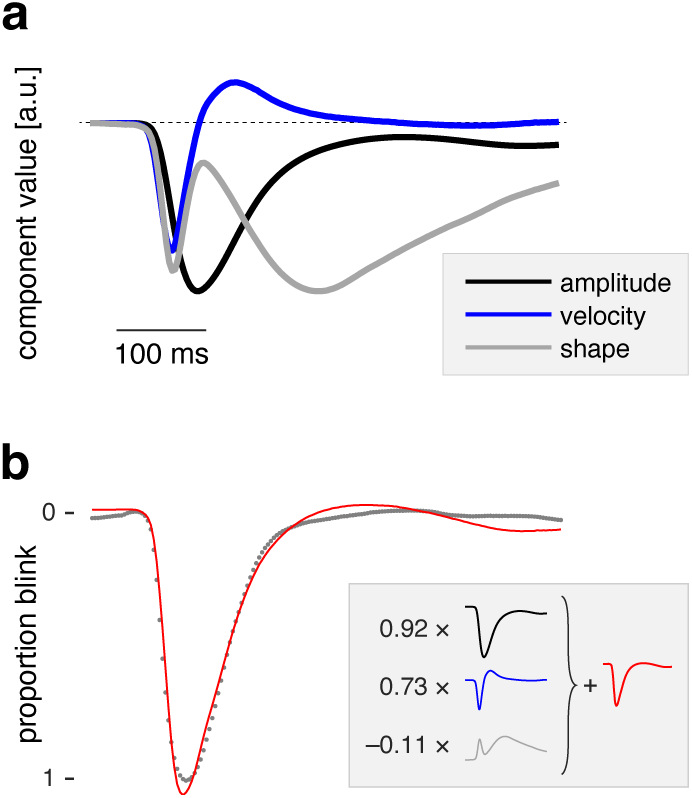
Temporal model components. **(a)** Lid position vectors over time were fit using a three-component temporal model. These components corresponded to the amplitude of the blink (black), the first-derivative of lid position and thus velocity (blue), and a third component that captured residual variation in shape (gray). **(b)** Example of the fit (red line) to a blink response (gray dots). Shown *inset* is the composition of this example fit, which is defined by the three coefficient values that scale the three temporal components. The sum of these scaled components provides the red fit line.

We examined the value of the fit coefficients averaged across participants for each stimulus intensity level. Consistent with our initial summary measures, the model coefficients demonstrate a systematic effect of air puff pressure upon blink response (Figure 4). The weight upon the first component (amplitude) increased with stimulus intensity, with a roughly linear increase in the proportion of closure of the palpebral fissure related to log changes in stimulus pressure (albeit with a plateau above 30 PSI). The weight upon the second component (velocity) also increased with stimulus intensity: log changes in stimulus intensity led to roughly linear increases in blink velocity, reducing the latency of the response by about 15 ms over the entire stimulus range. No systematic relationship between stimulus intensity and the third model component was seen. This last component was derived from the residual variation in the blink response that was not explained by variation in amplitude or velocity. The absence of a systematic relationship between stimulus pressure level and the coefficients of this third component argue against the existence of substantial, unmodeled variation in the response with pressure level that was not captured by the first two components.

**Figure 4.**
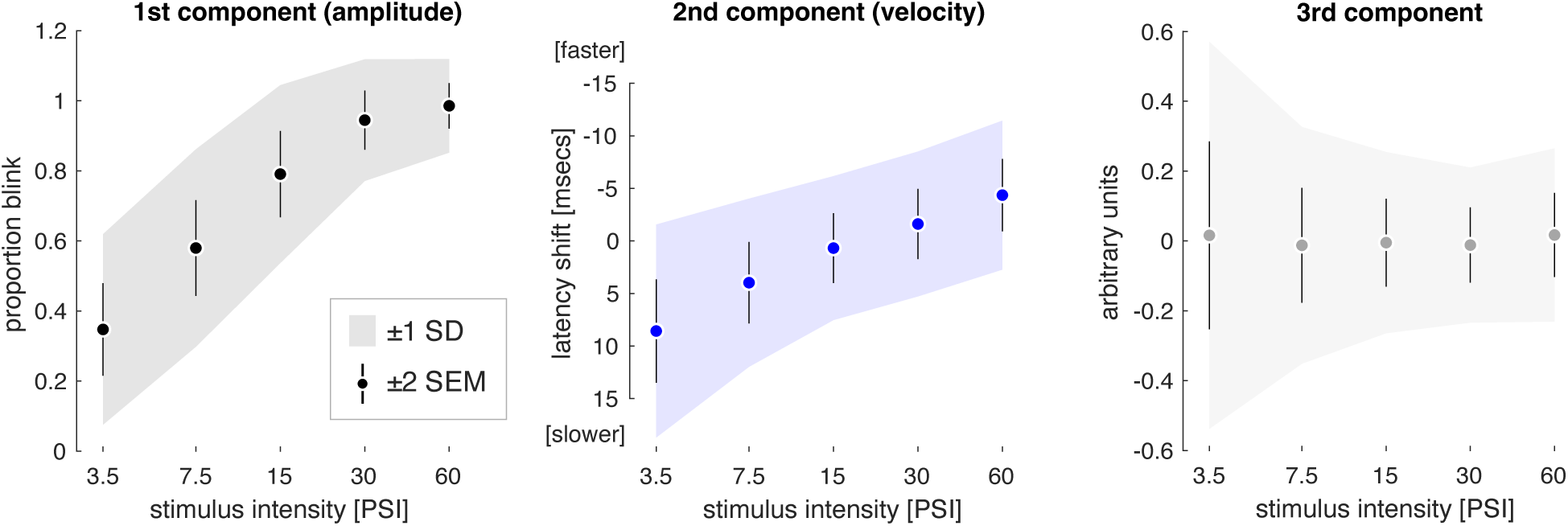
Coefficient values by stimulus intensity. Each plot presents the mean (across participant) coefficients for the three temporal (first (amplitude), second (velocity), and third) components as a function of stimulus intensity (log PSI). The shaded area is ±1 standard deviation of the mean across participants, and the error bars ±2 standard errors.

### 3.2 The contralateral blink response is subject to an increased stimulus threshold

The analyses up to this point combined the blink response measures from the eyelids ipsilateral and contralateral to the stimulus. We next tested if there are consistent differences in the sensitivity of the ipsilateral and contralateral response to mechanical somatosensory stimulation. We obtained the average amplitude coefficients across subjects for each stimulus level, separately for the ipsilateral and contralateral eye (Figure 5a). The measurements from each eye across pressure levels was then fit with a non-linear psychometric function (the Weibull cumulative distribution function). These fits provided a good account of the data and revealed that the intensity response function measured from the contralateral eye was shifted to the right as compared to the response from the ipsilateral eye. This property of the data indicates that higher intensity stimuli were required to evoke the same degree of lid closure in the contralateral eye as compared to the ipsilateral.

**Figure 5.**
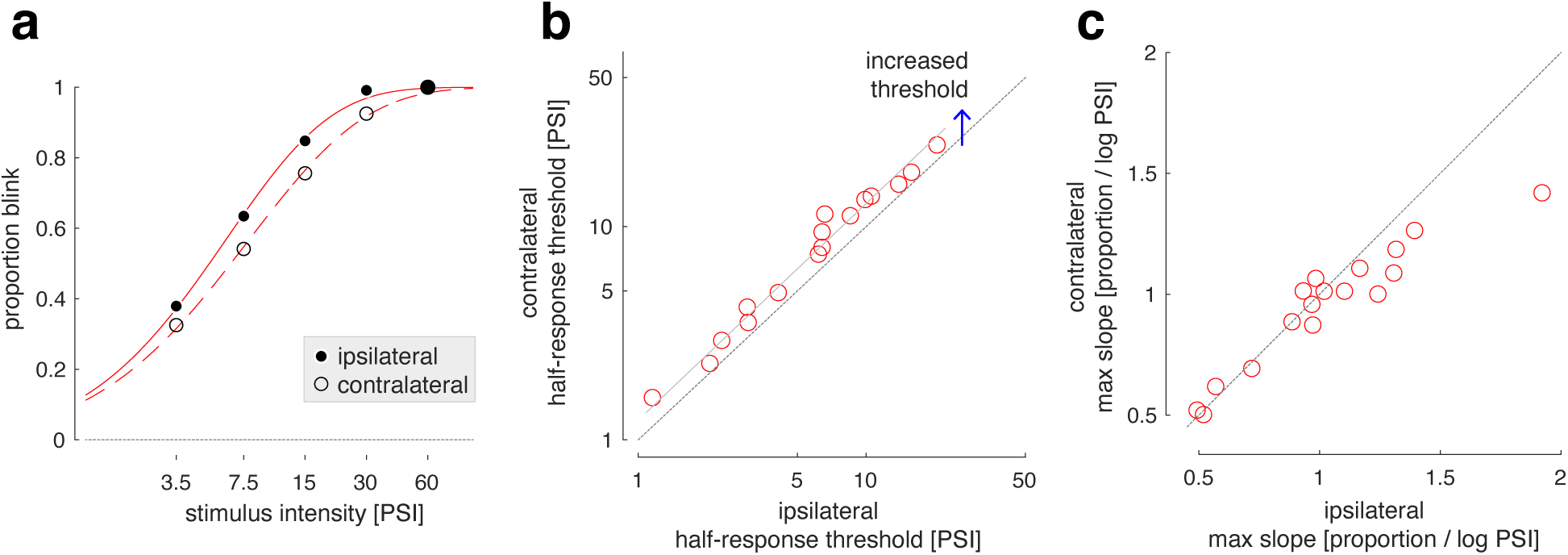
Ipsilateral versus contralateral blink responses. **(a)** The mean (across participant) amplitude coefficients as a function of stimulus intensity, obtained separately for the eyelid ipsilateral (filled circles) and contralateral (open circles) to the stimulus. Amplitude is expressed as the proportion of a full eyelid closure. The points are fit with Weibull cumulative distribution function (red lines). **(b)** The psychometric function was fit to the amplitude data for each participant and side (ipsilateral and contralateral), yielding two parameters (half-response threshold and maximum slope). Here, each plot point indicates the half-response threshold parameters measured for a particular participant from the ipsilateral and contralateral eyelid. A linear fit to the set of points (gray line) is parallel to and shifted upwards from the unit slope (dashed line), corresponding to an ∼0.1 log unit increased half-response threshold value measured from the contralateral as compared to the ipsilateral eyelid. **(c)** Each plot point indicates the maximum slope parameter value obtained from the ipsilateral and contralateral eyelid from each participant.

The Weibull function is under the control of two parameters that correspond to the stimulus intensity required to produce a 50% eye closure (the “half response threshold”) and the maximum slope of the change in eye closure with log changes in stimulus intensity. Figure 5b presents the half-response threshold values obtained for each participant from the ipsilateral and contralateral eyelid. While a wide range of values was seen, the plot points for all 17 participants were displaced ∼0.1 log units above the unit slope line. A paired t-test across participants of this measure between the contralateral and ipsilateral eye showed a significant difference (t[16 df] = 8.29, p = 3.499e-7). Thus, there was a highly reliable, ∼25% decrease in the threshold sensitivity of the response of the contralateral eyelid to somatosensory stimulation as compared to the ipsilateral eyelid.

Figure 5c presents the maximum slope of the psychometric function measured from the ipsilateral and contralateral eye for each subject. A wide range of values was again seen, although in this case the plot points did not systematically depart from the unit slope line, indicating a similar measurement from the two eyes. A paired t-test indicated that there was not a highly reliable difference in the slope value between the eyes (t[16 df] = 2.17, p=0.0453).

We examined if the timing of the response in the ipsilateral and contralateral eye differed. Across subjects and averaged across pressure levels, the velocity coefficient value was larger in the response from the contralateral eye as compared to the ipsilateral eye (t[16 df] = 3.83, p = 1.475e-03), corresponding to an approximately 4 ms relative delay in the response of the contralateral eye.

### 3.3 There is reliable individual variation in sensitivity to somatosensory stimulation

We next examined individual variation in the relationship between stimulus intensity and blink amplitude. Given that the difference between the ipsilateral and contralateral blink response was both small and consistent across individuals, we considered the blink responses averaged across the two eyelids. Figure 6a presents the fit of the Weibull function to the blink response amplitudes for each participant, ordered by the half-response threshold value obtained. For all participants, the psychometric function provided a good account of the effect of stimulus intensity upon blink amplitude, although the form of this relationship varied substantially. This variability can be seen both in the half-response threshold of the function (blue dotted lines), and in the maximum slope (steepness) of the fits.

**Figure 6.**
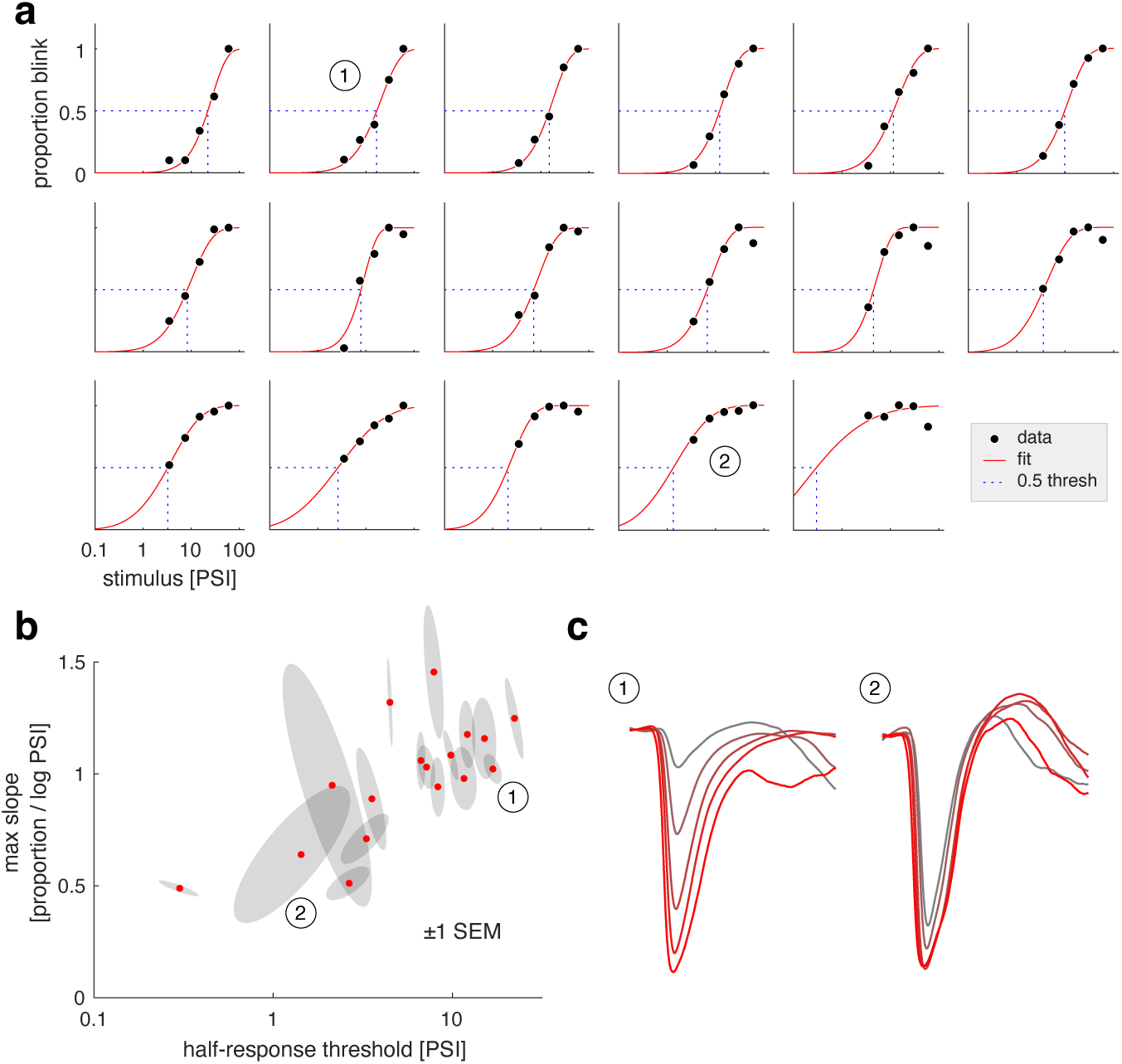
Individual differences in psychometric properties of blink amplitude. **(a)** Each sub-plot presents the amplitude coefficients for a participant as a function of stimulus intensity, averaged over acquisitions and the contralateral and ipsilateral eye. The fit of a Weibull cumulative distribution function is shown (red line), as is the half-response threshold value (dashed blue lines). Two example participants are identified with circled numbers, and these same participants are so indicated in subsequent panels. **(b)** The relationship between the half-response threshold and maximum slope parameters for each participant (red dots), surrounded by the bivariate normal distribution at one standard error of the mean obtained by bootstrap resampling across the 10 acquisitions (gray ellipses). **(c)** The average lid position plots at each stimulus intensity for two example participants (line color follows the convention of Figure 1).

The two parameters of the Weibull fit define for an individual the sensitivity of their blink response to varying somatosensory stimulation. Figure 6b plots the value of these two parameters for each participant, accompanied by the bivariate ellipse that indicates the standard error of the mean for the parameter. Differences between individuals were substantially larger than the measurement uncertainty. There was a positive correlation between half-response threshold and maximum slope across participants (r = 0.73). The underlying differences in blink response across stimulus levels is shown for two example participants in Figure 6c.

For roughly half of the participants, a decline in the amplitude of the blink was noted between the 30 and 60 PSI stimulus levels. Post-hoc examination of the data indicated that this phenomenon resulted from some subjects narrowing their palpebral fissure (i.e., squinting) during the set of eight trials with the highest stimulus intensity. This partial closing of the eye resulted in a smaller excursion of the lid in response to the stimulus.

### 3.4 Blink amplitude and velocity change with repeated stimulation

Finally, we examined how the properties of the blink response changed over the 8 trials that were presented in rapid succession within each acquisition. For each participant, we fit the 3-component temporal response model to the average response across stimulus intensity, acquisitions, and sides, separately for each of the 8 trials. Figure 7 shows that across trial number, there was a decrease in the amplitude of the blink response, a slight increase in velocity, and no change in the third component. These changes followed an exponential form, with the largest change in response coefficients being present between the first and second trial. We compared the R^2^ of the exponential model fit in the data to the R^2^ values obtained from fits to 10,000 permutations of the trial number order and found that the exponential form explains significant variance in the data for the amplitude (p=1.00e-4) and velocity (p=0.001) components, but not the third component (p=0.235).

**Figure 7.**
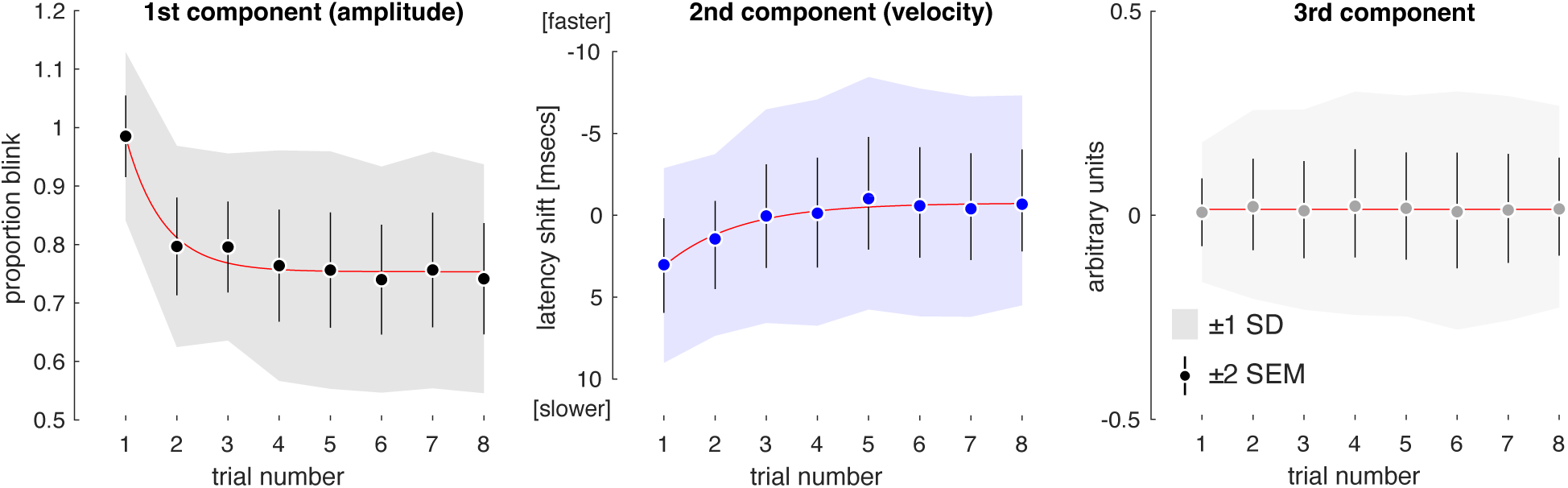
Coefficient values by trial number. Each plot presents the mean (across participant) coefficients for the three temporal (first (amplitude), second (velocity), and third) components as a function of trial number. The shaded area is ±1 standard deviation of the mean across participants, and the error bars ±2 standard errors. The fit of an exponential function is shown (red line).

### 3.5 The auditory component of the stimulus alters the later stages of the blink response

The delivery of the air puff was accompanied by an audible hiss. We measured sound pressure at the ear of the experimental subject and found that it grows steadily with the stimulus intensity (Figure S1b), becoming quite loud (83 dB) at the highest stimulus level. To examine the effect of the auditory component, we conducted a control experiment (n = 10) in which we measured the blink elicited by the 30 PSI stimulus under conditions in which the auditory component was, or was not, masked by a noise stimulus played over sound-cancelling headphones (Figure S2). The amplitude and latency of blink responses were very similar up to ∼130 ms for the measurements with and without an auditory cue. Beyond 130 ms, the eyelid returned to the open position faster when accompanied by an auditory cue, and in some subjects the palpebral fissure opened wider than in its initial position.

## 4. Discussion

We have developed a measure of parametric variation in the somatosensory blink reflex and have shown that there are substantial individual differences in the sensitivity of this response to changes in stimulus intensity. There have been studies of parametric variation in auditory (Peak, 1932) and visual stimulation (Bixler, Bartlett, & Lansing, 1967) which generally show an increase in amplitude and a decrease in blink latency with increasing stimulus intensity. We are unaware of prior work that has quantified how the form and timing of the human blink response changes with mechanical stimulus intensity, or how these changes covary within or across people.

Prior measurements of the somatosensory blink reflex have been conducted primarily using electrical stimulation and measurement of an EMG response; here we use mechanical stimulation and a measure of the physical movement of the eyelid. A strength of our measurement is that we have studied the full, biological response (eyelid closure) to an ecologically valid stimulus (mechanical stimulation of the somatosensory surface of the eye and periorbital skin). This measure may be more sensitive to pathologic abnormalities in the somatosensory blink reflex, albeit with less etiologic specificity. An additional advantage of our measure is that it may be readily used within the magnetic resonance imaging environment, thus supporting studies of the neural correlates of trigeminal stimulation(Lee, et al., 2017).

An important caveat to our results is that the delivery of the air puff was accompanied by an audible hiss. The auditory component of our stimulus arose at the nozzle adjacent to the eye, and thus the somatosensory stimulus and auditory cue were essentially simultaneous. An acoustic cue that precedes an air-puff stimulus (e.g., by 70 ms) inhibits the corneal blink response (Krauter, Leonard, & Ison, 1973), but can transition to an excitatory effect when presented simultaneously (Flaten & Blumenthal, 1998). In a control experiment, we find that the auditory component of the stimulus has minimal impact upon the initial (<130 ms) time period of the blink response but is associated with a more rapid re-opening of the eye at later time points. As our study focused upon the amplitude and latency properties within the early response regime, we think it likely that auditory component had a relatively small impact upon our results. Reassuring in this regard is the observation that the increase in stimulus intensity at the lowest stimulus levels (3.5 and 7.5 PSI) produced a proportional change in blink response, despite these two stimuli having nearly equivalent auditory components (Figure S1b). Nonetheless, future studies would be improved by masking the auditory component.

Our approach is also limited in separating the individual response components observed in EMG (e.g., R1, R2, R3). In primates, the early component (R1) is a small, ipsilateral, oligosynaptic reflex, while the later, larger component (R2) is a bilateral, polysynaptic response (Vollono, 2021). It is the R2 that corresponds with the actual blink of the eyelids (Rossi, Risaliti, & Rossi, 1989). In rabbits, an air puff applied to the cornea elicits a monophasic blink that incorporates the R1 and R2 EMG responses (Gruart, Schreurs, del Toro, & Delgado-Garcia, 2000); our measurements likely reflect this combined response as well. A third, still later EMG component (R3; latency of 75 to 90 ms) has also been observed. This was initially reported in response to noxious stimuli (Rossi, Risaliti, & Rossi, 1989), but later observations found it present independent of nociception (Ellrich, Katsarava, Przywara, & Kaube, 2001; Tellez, Axelrod, & Kaufmann, 2009) and may represent a startle reflex (Brown, et al., 1991). After 200 ms, we also observe in some subjects widening of the palpebral fissure as an effect of the auditory component of the stimulus (Figure S2). We speculate that this is the retraction of the upper lid that has been identified as a marker of the emotion of “surprise” (Ekman, 1978). We have been unable to find a clear, prior measurement of the time course of this lid retraction in people, and we note that orbicular oculi EMG would not reflect this signal from the levator palpebrae superioris.

We have shown that blink amplitude is well described by a two-parameter psychometric function of the log of trigeminal stimulation intensity. The psychometric function is fully described by the threshold and slope parameters. We find a systematic difference between the threshold parameters measured from the eyelid ipsilateral and contralateral to the stimulus. That is, a higher intensity stimulus is required to evoke a given amount of lid movement in the contralateral eye as compared to the ipsilateral eye. Consistent with this, prior studies have found that the blink response contralateral to an air puff to be smaller and slower (VanderWerf, Brassinga, Reits, Aramideh, & Ongerboer de Visser, 2003). A notable feature of our data is that, despite marked individual variation in overall threshold and slope, the threshold difference between the ipsilateral and contralateral blink response function is nearly identical across individuals (Figure 5b). This suggests that the increased contralateral threshold arises from a common neural mechanism across individuals, operating downstream from the source of individual variation.

We find that there are substantial differences amongst participants in the overall sensitivity of the somatosensory blink reflex, as indexed both by half-response threshold and the slope of the response function. There is a relatively strong correlation across individuals between these two parameters, suggesting that people vary in a primary sensitivity axis. The site of this variation could be present anywhere along the reflex pathway, and a goal for future work is to identify behavioral or perceptual correlates of this variation, and the neural source of the modulatory effect.

In other sensory domains, measurement of a stimulus-response function offers a fundamental characterization of sensitivity that can have clinical importance (Balcer, et al., 2000). For the blink reflex, measurement of the response function may reveal alterations in the sensitivity of the trigeminal system in clinical disorders. An intriguing target is migraine, for which potentiation of pain signals from trigeminal afferents is a cardinal feature. Despite the apparent association between the trigeminal system and migraine, there is limited consensus regarding what (if any) alterations of somatic sensory processing accompany headache disorders (Coppola & Chen, 2021). EMG measures of the trigeminal blink response have been extensively studied in migraine, but with conflicting results regarding if an alteration in response is present (Vollono, 2021). It may be the case that these studies, performed at a single stimulus intensity level, have missed differences that would be revealed in the full trigeminal sensitivity function.

## 5. Conclusion

We find that parametric variation in the intensity of mechanical stimulation of the trigeminal nerve produces a corresponding, graded response in the velocity and amplitude of the blink response. The function that relates stimulus intensity to blink amplitude has a common form but different sensitivity parameters across individuals. We view this as a promising framework for investigating group and individual differences in the sensitivity of the trigeminal system.

## Acknowledgements

None

## Author Contributions

Conceptualization: E.K., V.Y.B., and G.K.A. Data curation: G.K.A. Formal analysis: E.B.H. and G.K.A. Funding acquisition: V.Y.B. and G.K.A. Investigation: E.B.H. and D.P.G. Methodology: E.B.H. and G.K.A. Project administration: E.B.H. Resources: D.P.G. Supervision: D.P.G. and G.K.A. Visualization: E.B.H. and G.K.A. Writing - original draft: E.K., E.B.H., and G.K.A. Writing - review & editing: E.K., E.B.H., D.P.G., V.Y.B., and G.K.A.

## Data availability statement

The available raw data are of the form of lid position vectors recorded for every trial. These data, and the code needed to reproduce the analysis and figures presented here, are publicly available: https://github.com/gkaguirrelab/Kaiser_2023_Psychophysiology

## Supplementary materials

### Physical properties of the stimulus

#### Spatial extent

The air puff is delivered via a nozzle that is 2.3 mm in diameter, positioned ∼20 mm from the lateral canthus of the subject. The stimulus approximates a compact, isothermal, free-air jet. At the point of impact with the face, the air jet is greater than 8 nozzle diameters distant from the nozzle and thus is in the region of fully developed turbulence (Zhivov et. al., 2020). Within this region—and accounting for the additional 6 mm distance to the origin point of the modeled conical air jet—the spatial velocity profile of the stimulus is Gaussian, with width at plus / minus two standard deviations of roughly:

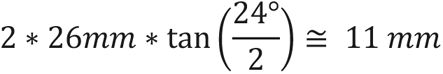

Figure S1a illustrates this geometry and the surface area around the lateral canthus that receives the stimulus.

#### Maximum sound pressure amplitude

We measured the acoustic pressure amplitude of the stimulus by placing a sound level meter (Tadeto Digital Sound Level Meter Portable SPL) at the position of the right ear. The maximum pressure amplitude was obtained across 8 presentations of each of the 5 stimulus intensity levels. We also measured the ambient level of the environment. Figure S1b presents these data and shows that the amplitude of the highest intensity stimulus was 83.4 dB.

#### Temporal profile

We placed a microphone (MAONO USB Lavalier Microphone) at the position of the right lateral canthus and measured the average temporal profile of the stimulus across four air-puff trials at 8000 Hz. Figure S1c presents the average rectified recording for each of the stimulus intensity levels. The arrival of the stimulus is marked by a sharp onset of acoustic energy; there is likely clipping of the absolute value of this measurement. Then follows a 100 ms period of a steady stream of air flow, with the amplitude of this component increasing across stimulus levels. Finally, there is again a transient increase in acoustic energy at stimulus offset.

**Figure S1.**
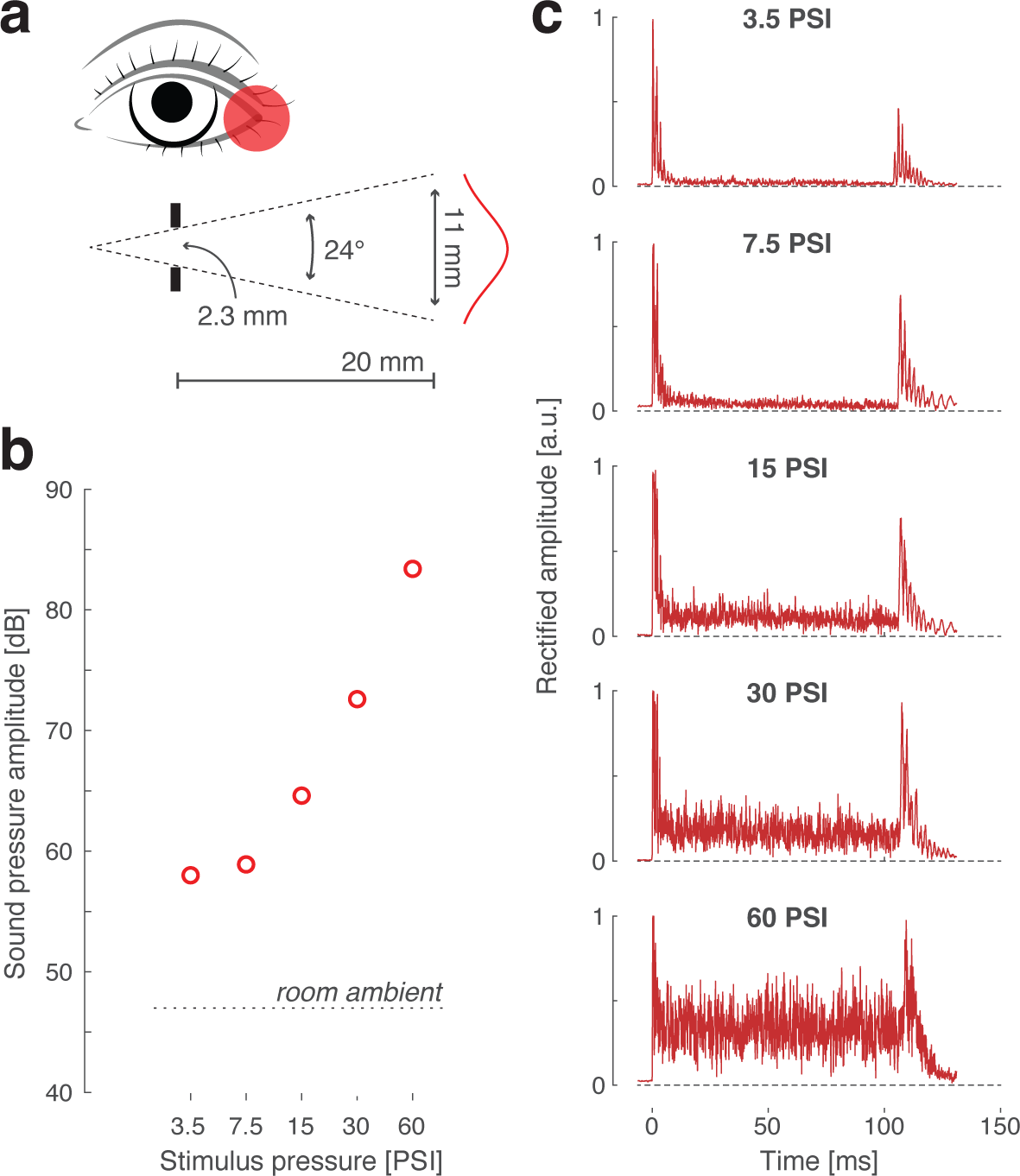
Physical properties of the stimulus. a) The air puff was delivered from a nozzle 2.3 mm in diameter that was positioned ∼20 mm from the lateral canthus of the participant. The velocity profile of a free air jet with this geometry is illustrated. b) The maximum recorded pressure amplitude of each stimulus intensity level at the position of the ear of the participant. The dashed line is the ambient sound pressure amplitude of the room in the absence of the stimulus. c) The temporal profile of the rectified amplitude of the recorded sound of the stimulus at each pressure level.

### Effect of the acoustic component of the stimulus upon the blink response

We measured the blink response to the 30 PSI stimulus in 10 participants not involved in the original data collection. A total of 7 acquisitions were performed, and each acquisition included 8 air puff trials. The data from the first acquisition were discarded. During acquisitions 2, 3, and 5, the participants wore noise-canceling headphones, which additionally played a continuous “brown noise” sound at near maximum volume. For the other acquisitions, the participant did not wear headphones. Participants reported that the auditory component of the air puff stimulus was almost or completely eliminated by the headphones and continuous noise.

The first air puff trial of each of the 6 retained acquisitions was discarded. For each participant, we obtained the average ipsilateral and contralateral blink response across trials and acquisitions, separately for the acquisitions with and without masking the stimulus’s acoustic properties. Blink amplitude was expressed as a proportion of lid closure.

**Figure S2.**
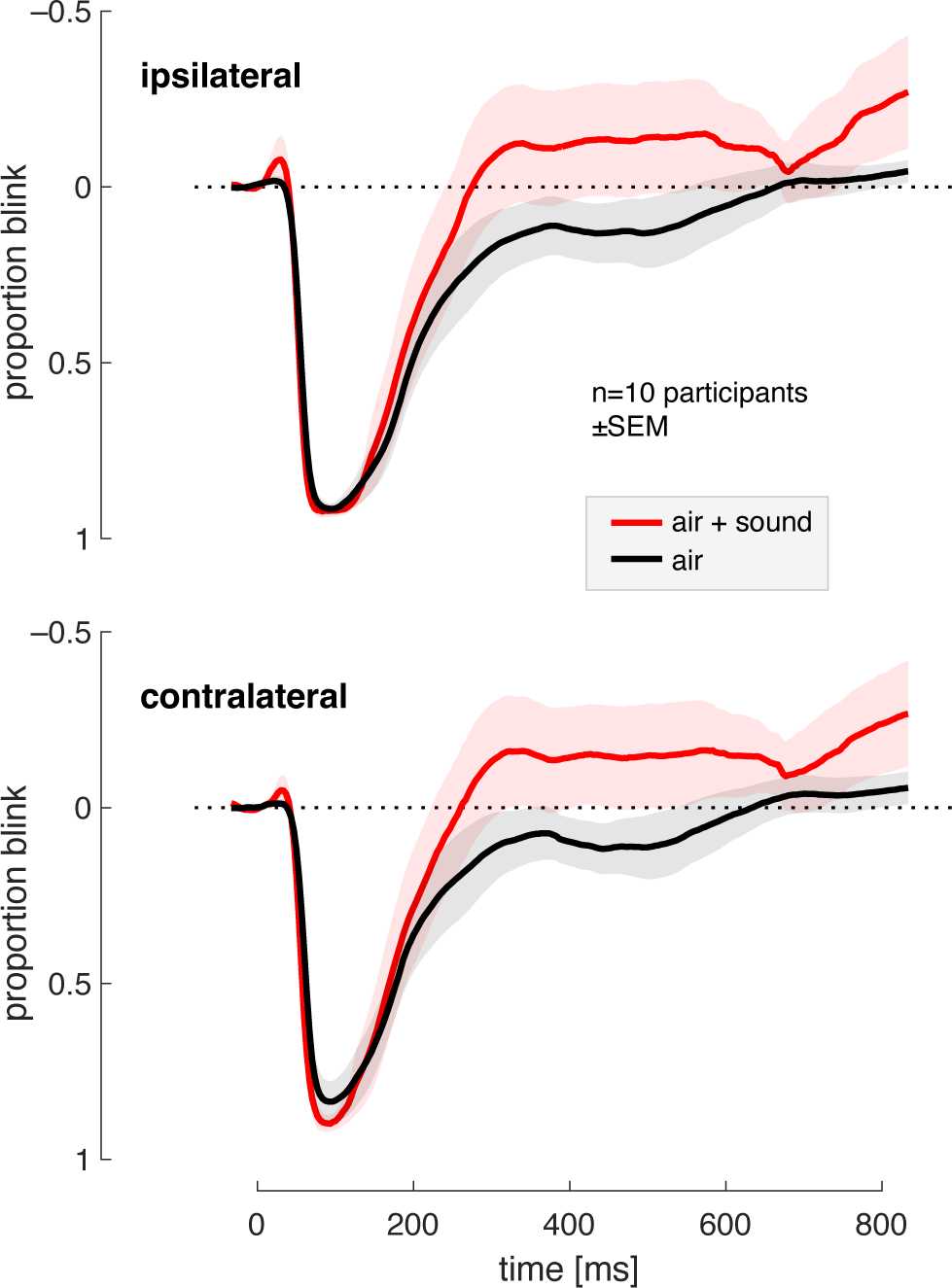
Average, across subject blink responses from the eye ipsilateral (*top*) and contralateral (*bottom*) to a 30 PSI stimulus. Separate responses were recorded for the stimulus accompanied by the auditory component (“air + sound”) and when the stimulus was the air puff alone (“air”). Shaded areas indicate ±1 SEM across ten participants.

The average responses across participants are presented in Figure S2. The amplitude and latency of blink responses are very similar, up to ∼130 ms for the measurements with and without auditory cues. Beyond 130 ms, the eyelid returns to the open position faster when accompanied by an auditory cue, and in some subjects, the palpebral fissure opens wider than in its initial position.

